# UpSetR: An R Package for the Visualization of Intersecting Sets and their Properties

**DOI:** 10.1101/120600

**Authors:** Jake R Conway, Alexander Lex, Nils Gehlenborg

## Abstract

Venn and Euler diagrams are a popular yet inadequate solution for quantitative visualization of set intersections. A scalable alternative to Venn and Euler diagrams for visualizing intersecting sets and their properties is needed. We developed UpSetR, an open source R package that employs a scalable matrix-based visualization to show intersections of sets, their size, and other properties. UpSetR is available at *https://cran.r-project.org/package=UpSetR* and released under the MIT License. A Shiny app is available at *https://gehlenborglab.shinyapps.io/upsetr/*.

## 1 Introduction

The visualization of sets and their intersections is a common challenge for researchers who are dealing with biological and biomedical data. For example, a researcher might need to compare multiple algorithms that identify single nucleotide polymorphisms (Xu, F., et al., 2012, Supplementary Figure 1) or show orthologs of genes in newly sequenced species across genomes of related species (D’Hont, A., et al., 2012, Supplementary Figure 2). Although many alternative set visualization techniques exist (Alsallakh, B., et al., 2014), such data is typically visualized using Venn and Euler diagrams. Such diagrams can be generated with R packages such as *venneuler* (Wilkinson, 2012) and *VennDiagram* (Chen and Boutros, 2011). These closely related techniques have well known shortcomings, as they are hard to generate for more than a small numbers of sets. The visual representation of intersection size by irregularly shaped and unaligned areas makes it hard to answer essential questions such as “What is the biggest intersection?” or “Is intersection *X* larger than intersection *Y*?” (Cleveland and McGill, 1984).

## 2 Methods

Here we present an R package named “UpSetR” based on the “UpSet” technique (Lex, A. et al., 2014) that employs a matrix-based layout to show intersections of sets and their sizes. It is implemented using ggplot2 (Wickham, 2009) and allows data analysts to easily generate generate UpSet plots for their own data. UpSetR support three input formats: (1) a table in which the rows represent elements and columns include set assignments and additional attributes; (2) sets of elements names; and (3) an expression describing the size of the set intersections as introduced by the *venneuler* package (Wilkinson, 2012). UpSetR provides support for the visualization of attributes associated with the elements contained in the sets, enabling researchers to explore and characterize the intersections. UpSetR differs from the original UpSet technique as it is optimized for static plots and for integration into typical bioinformatics workflows. We also provide a Shiny app that allows researchers to create publication-quality UpSet plots directly in a web browser.

UpSetR visualizes intersections of sets as a matrix in which the rows represent the sets and the columns represent their intersections (Figure 1 and Supplementary Figures 1 and 2 for comparisons of Venn and Euler diagrams with UpSetR plots). For each set that is part of a given intersection, a black filled circle is placed in the corresponding matrix cell. If a set is not part of the intersection, a light gray circle is shown. A vertical black line connects the topmost black circle with the bottommost black circle in each column to emphasize the column-based relationships. The size of the intersections is shown as a bar chart placed on top of the matrix so that each column lines up with exactly one bar. A second bar chart showing the size of the each set is shown to the left of the matrix.

**Figure 1:**
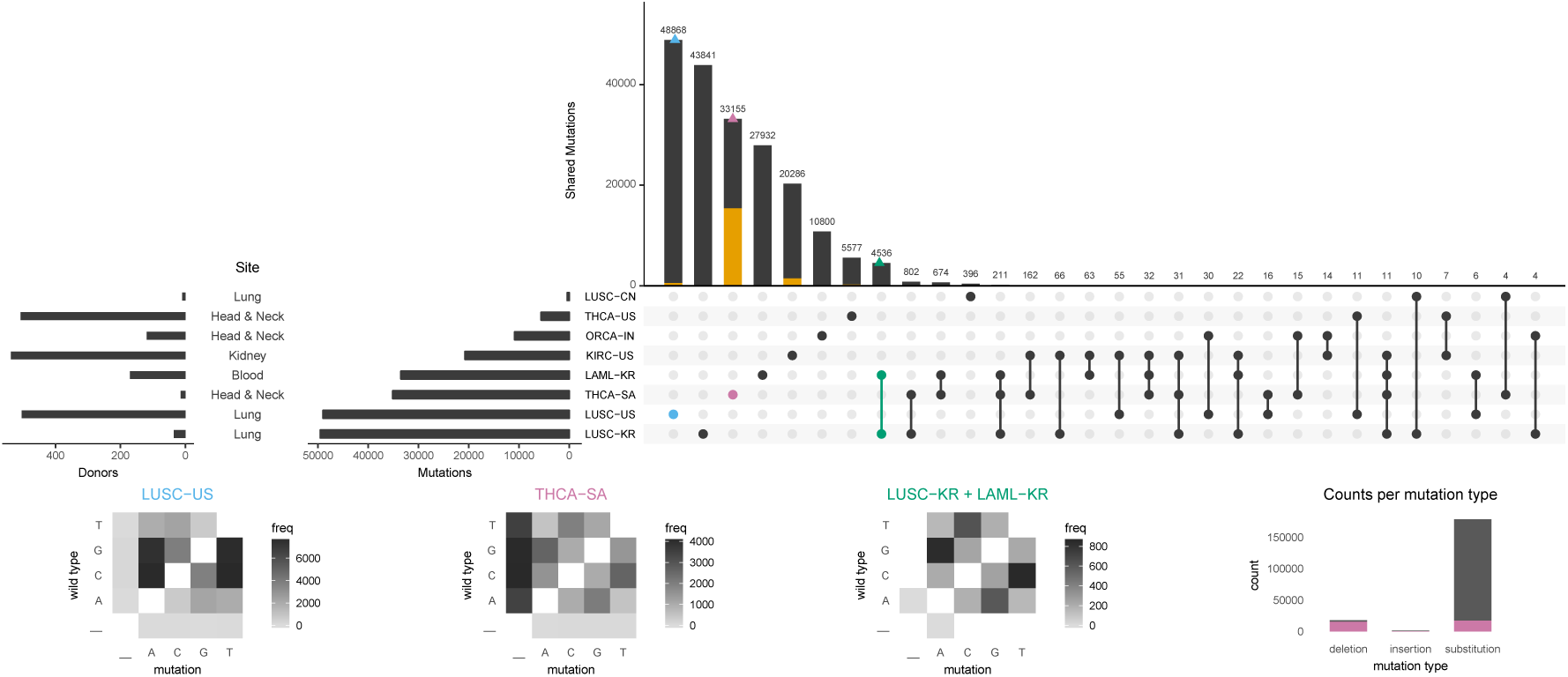
An UpSetR plot of variants across eight ICGC cancer studies with three intersection queries, one element query, four attribute plots, and two set metadata plots. The three intersection queries are the one-way intersections of LUSC-US (blue) and THCA-SA (purple), and the two-way intersection of LUSC-KR and LAML-KR (green). The element query (yellow) selects mutations classified as deletions. Three custom transition/transversion plots display the relative frequency of substitution events for the intersection queries. The bar plot attribute plot displays the contribution of variants unique to the THCA-SA cohort (purple) to each mutation type. Set metadata is plotted to the left of the set size bar. charts.

## 3 Usage Scenario

To illustrate the utility and features of UpSetR, we retrieved variant calls for eight cancer studies from from the ICGC Data Portal (see Supplementary Information). Each cancer study represents a set and each variant represents an element that is contained in one or more sets (Supplementary Figure 3). UpSetR supports queries on the data to highlight features. *Intersection queries* can be used to select subsets of elements in the data set defined by an intersection. Queries are assigned a unique color and their results are plotted on top of the intersection size bar chart. For example, this can be used to select elements in particular intersections (Supplementary Figure 4). Additionally, UpSetR supports queries for the selection of elements based on attributes associated with the elements in the sets. Attributes can be numerical, Boolean, or categorical. In our example, element attributes are chromosome, genomic location, and variant type (deletion, insertion, substitution) associated with each variant. UpSetR *element queries* select elements across intersections and sets based on particular attribute values. Basic built-in queries can be extended to arbitrarily complex queries by providing a custom query function that operates on any combination of attributes. Element queries can be used to select variants of a particular type, such as deletions, and to view them across intersections (Supplementary Figure 5).

UpSetR provides integration of additional *attribute plots* that visualize attributes of elements selected by an intersection or element query. Support for scatter plots and histograms is built into UpSetR. Additional plot types can be integrated by providing in a function that returns a *ggplot* object to visualize the data. When attribute or intersection queries are applied, query results can also be overlaid on attribute plots in addition to the intersection size bar plot. Figure 1 demonstrates how these features, including the visualization of metadata about the sets, can be combined into a plot that among other issues, reveals a notable over-representation of unique deletions among the variants in one of the eight cancer studies retrieved from the ICGC data portal.

## 4 Conclusion

UpSetR is a highly customizable tool for data exploration and generation of set visualizations. By making UpSetR compatible with the input formats of existing popular Venn and Euler diagram packages and by offering a Shiny web interface, we incentivize use of UpSet diagrams and enable users without programming skills to generate effective set visualizations. Through its seamless integration with ggplot2 and its ability to apply virtually any query, it is possible to customize and explore data in ways not supported by any other set visualization package. In addition, the integration of UpSetR with ggplot2 allows developers to extend UpSetR for use in their own software packages.

## Acknowledgements

The authors would like to thank Megan Paul for her contributions to the project. This work was funded through awards by the National Institutes of Health (R00HG007583, U54HG007963, U01CQ198935).

